# A decisional space account of saccadic reaction times towards personally familiar faces

**DOI:** 10.1101/292656

**Authors:** Meike Ramon, Nayla Sokhn, Roberto Caldara

## Abstract

Manual and saccadic reaction times (SRTs) have been used to determine the minimum time required for different types of visual categorizations. Such studies have demonstrated that faces can be detected within natural scenes within as little as 100ms (Crouzet, Kirchner & Thorpe, 2010), while increasingly complex decisions require longer processing times (Besson, Barragan-Jason, Thorpe, Fabre-Thorpe, Puma et al., 2017). Following the notion that facial representations stored in memory facilitate perceptual processing (Ramon & Gobbini, 2018), a recent study reported 180ms as the fastest speed at which “familiar face detection” based on expressed choice saccades (Visconti di Ollegio Castello & Gobbini, 2015). At first glance, these findings seem incompatible with the earliest *neural* markers of familiarity reported in electrophysiological studies (Barragan-Jason, Cauchoix & Barbeau, 2015; Caharel, Ramon & Rossion, 2014; Huang, Wu, Hu, Wang, Ding & Qu et al., 2017), which should temporally *precede* any overtly observed behavioral (oculomotor or manual) categorization. Here, we reason that this apparent discrepancy could be accounted for in terms of *decisional space* constraints, which modulate both manual RTs observed for different levels of visual processing (Besson et al., 2017), as well as saccadic RTs (SRTs) in both healthy observers and neurological patients (Ramon, in press; Ramon, Sokhn, Lao & Caldara, in press). In the present study, over 70 observers completed three different SRT experiments in which decisional space was manipulated through task demands and stimulus probability. Subjects performed a gender categorization task, or one of two familiar face “recognition” tasks, which differed with respect to the number of personally familiar identities presented (3 vs. 7). We observe an inverse relationship between visual categorization proficiency and decisional space. Observers were most accurate for categorization of gender, which could be achieved in as little as 140ms. Categorization of highly predictable targets was more error-prone and required an additional ~100ms processing time. Our findings add to increasing evidence that pre-activation of identity-information can modulate early visual processing in a top-down manner. They also emphasize the importance of considering procedural aspects as well as terminology when aiming to characterize cognitive processes.

## Introduction

The human visual system can rapidly perform numerous tasks across large variations in stimulus input, and with extremely high efficiency. For instance, categorization of animals presented in images of natural scenes presented for as little as 20ms can be reliably achieved in under 300 milliseconds (ms) (e.g., Thorpe, Fize & Marlot, 1996; Rousselet, Fabre-Thorpe & Thorpe, 2002; Macé, Thorpe, Fabre-Thorpe, 2005). Among animate objects, human faces appear to have a special status: they can be rapidly categorized (detected within natural scenes) within as little as 100-110ms (Crouzet, Kirchner & Thorpe, 2010). Such rapid categorical responses have been used to study the characteristics of visual processing. Specifically, the time required to perform such accurate visual categorizations provide a valuable source of information that can be levered to constrain theories of visual processing (for a review see e.g., Fabre-Thorpe, 2011).

In attempting to maximize the precision estimates of visual processing speed, different paradigms have been developed (for a direct comparison of paradigms, see Bacon-Macê, Kirchner, Fabre-Thorpe & Thorpe, 2007). Estimates of visual categorizations have been derived from verification and naming tasks, which typically involve longer reaction times (RTs) (Tanaka & Taylor, 1991) compared to manual forced choice paradigms, where subjects press (or release) one of two buttons to distinguish between stimulus categories (e.g., Hasbroucq, Mouret, Seal, & Akamatsu, 1995). To reduce response times, more sensitive Go/no-go paradigms have been developed that require subjects to respond to a predefined target category by finger-lift from an infra-red sensor (e.g., Thorpe et al., 1996; VanRullen & Thorpe 2001; Rousselet et al., 2002; Bacon-Macé, Macé, Fabre-Thorpe & Thorpe, 2005). Finally, other paradigms exploiting the very rapid responses from oculo-motor, as compared to manual, effectors have been developed to provide a more precise description of the lower bound of speeded categorizations (Kirchner & Thorpe, 2006; Crouzet, Joubert, Thorpe & Fabre-Thorpe, 2009; Crouzet et al., 2010). Aside from deriving mean or median RTs, these paradigms have aimed to determine the (manual or saccadic) *minimum* reaction time (minRT). The minRT is defined as the first time-bin for which correct responses significantly outnumber incorrect ones (Fabre-Thorpe, Richard & Thorpe, 1998; VanRullen & Thorpe, 2001), and is considered to reflect the minimal processing time required for *reliable* responses (Rousselet, Macè & Fabre-Thorpe, 2003).

Visual processing speed of faces is, however, not only determined by the paradigm opted for, but also depends on the nature of visual categorization performed (Barragan-Jason, Lachat & Barbeau, 2012), which impacts upon information diagnosticity (Schyns, 1998), as does prior familiarity (Ramon, Caharel & Rossion, 2011; for a review see Ramon & Gobbini, 2018). For example, superordinate human vs. animal decisions can be performed manually with high precision (98%) in as little as 285ms, while famous vs. unfamiliar manual decisions are more error-prone and slower (75%, 468ms); gender categorization on the other hand can be performed with high fidelity and intermediate minRTs (94%; ~310ms) (Barragan-Jason et al., 2012). Such findings have led to the suggestion that visual processing involves initial feed-forward propagation of activity through occipito-temporal regions (Thorpe et al., 1996), providing coarse magnocellular representations, which require additional processing and refinement before higher-level categorization can be achieved (Fabre-Thorpe, Delorme, Marlot & Thorpe, 2001; Macé, Joubert, Nespoulous & Fabre-Thorpe, 2009; Fabre-Thorpe, 2011). In short, this evidence suggests that face detection (i.e., ascertaining the presence or location of a face) occurs before establishing the familiarity of a face.

The idea that face detection precedes familiarity recognition is also supported by a recent carefully conducted study reported by Besson and colleagues (Besson, Barragan-Jason, Thorpe, Fabre-Thorpe, Puma et al., 2017). This study involved a Speed and Accuracy Boosting procedure – a variant of the manual go/no-go paradigm that includes a response deadline and performance feedback designed to constrain subjects to use their fastest strategy. Using this procedure in combination with extremely large stimulus sets to avoid image repetitions, the authors reported manual minRTs that differed as a function of task. Both human (vs. animal) face recognition and individual face recognition (i.e., searching for a single predefined target identity) were performed in a highly accurate and extremely fast manner (min RTs: ~240ms and ~260ms). On the other hand, deciding whether a face belonged to a large pool of famous individuals (“famous face recognition”) was more error prone and required more time (~380ms), in line with findings from speeded go/no-go paradigms with personally familiar faces (Ramon et al., 2011). Together these findings support the idea that 380ms may represent the lower bound for speeded manual familiarity decisions that occur ca. 180ms after the earliest neural marker of familiarity (e.g., Caharel, Ramon & Rossion, 2014; Barragan-Jason, Cauchoix & Barbeau, 2015; Huang, Wu, Hu, Wang, Ding & Qu et al., 2017).

One recent study seems incompatible with these independent lines of evidence. Measuring the minimum speed of choice saccades expressed towards personally familiar (PF) faces presented simultaneously with unfamiliar face (UF) distractors, Visconti di Oleggio Castello and Gobbini (2015) reported remarkably fast personally familiar face “detection” in 180ms. Naturally, overtly expressed saccadic behavior would require *prior* successful neural discrimination. However, the minimum saccadic reaction times (minSRTs) reported by Visconti di Oleggio Castello and Gobbini (2015) concur with or precede the earliest familiarity-dependent, differential electrophysiological response to faces reported to date (~140-200ms; e.g. Caharel et al., 2014; Barragan-Jason et al., 2015).

We reasoned that procedural aspects could account for these apparently discrepant findings. On the surface, Visconti di Oleggio Castello and Gobbini’s (2015) task seems comparable to the famous face recognition reported by Besson et al. (2017) and Ramon et al. (2011): all required categorical, i.e. binomial decisions of familiarity stimuli on a trial-by-trial basis. However, an important (and in our opinion neglected) aspect concerns the *number of stimuli* comprising, and thereby *constraining observers decisional space*, which can dramatically affect observers categorization proficiency (Besson et al., 2017; Ramon, Sokhn & Caldara, 2017; Ramon, Sokhn, Lao & Caldara, in press). A fundamental *difference* between the aforementioned electrophysiological and behavioral studies concerns the number of identities based on which observers’ performed their binary decisions. Visconti di Oleggio Castello and Gobbini’s (2015) group of seven observers performed choice saccades towards a total of *three* PF target identities. Despite the several hundreds of repetitions of each stimulus over the course of their experiment^1^, their observers accomplished this task with modest performance accuracy (average: 62%; range: 49-69%), in as little as 180m^2^. They concluded that “detectors for diagnostic features associated with overlearned familiar faces” (p.2), “allow rapid detection that precedes explicit recognition of identity” (p.1). Using a comparable SRT paradigm, Mathey and colleagues (Mathey, Besson, Barragan-Jason, Garderes, Barbeau & Thorpe 2012) reported even shorter minSRTs of <140ms, along with comparatively higher performance accuracy (range: 60-75%). Interestingly, their observers performed an “individual face recognition” task, in which a *single* famous identity served as the sole target. Based on their findings, the authors suggested that “information about identity can impact behavior much faster than had been previously suspected” (p.166).

Thus, both studies involved SRT target search paradigms (Visconti di Oleggio Castello & Gobbini, 2015; Mathey et al., 2012) and reported extremely rapid (S)RTs executed towards familiar faces. Comparing the reported results, it is appears that the decrease the number of target items (3 targets vs. 1 target) could account for the observed performance difference (lower vs. higher accuracy; slower vs. shorter minSRTs), which results from an effective narrowing of observers’ decisional space. Interestingly, both studies draw vastly different conclusions based on their convergent findings. While Mathey et al. (2012) suggest that “information about identity can impact behavior much faster than had been previously suspected” (p.166), Visconti di Oleggio Castello and Gobbini (2015) concluded that “detection […] precedes explicit recognition of identity” (Visconti di Oleggio Castello & Gobbini, 2015; p.1). In our opinion, the latter conclusion is not warranted, as their study involved *1-of-3 target search task* (cf. Besson et al., 2017; Ramon et al., 2017), which involves different task demands than a detection task. This stresses the importance of considering procedural aspects in concert with the appropriateness of the adopted terminology (cf. Ramon, in press).

Expanding on these previous findings, the present study sought to determine the lower boundary for fast visual categorization of personally familiar faces, and ascertain the effect of stimulus expectations on processing speed. Observers completed three different experiments involving binary decisions by performing two alternative forced-choice saccades: a gender categorization task, as well as two familiarity decision tasks (comparable to those reported previously; Ramon et al., 2011; Visconti di Oleggigo Castello & Gobbini, 2015; Besson et al., 2017; Ramon et al., 2017). The latter two tasks differed in terms of the number of personally (un)familiar identities presented – observers performed a 1-of-*n*-target search task. This effectively manipulates observers’ expectation: both familiarity decision tasks differed in that their *decisional space* was either comparatively *broad* or *narrow*.

We anticipated that performance would be highest and SRTs fastest in the gender categorization task, which did not require identity individuation, as subjects were instructed to saccade towards target stimuli belonging to one of two categories (female faces). Although the familiarity tasks also involved binary decisions (saccades towards PF faces), we anticipated that performance would depend on the target / distractor category set size used. Specifically, we expected higher performance and faster SRTs for a narrow decisional space – when target identities were highly predictable, given a smaller sample of PF faces to detect. In keeping with Besson et al (2017), but contrary to Visconti di Oleggio Castello and Gobbini (2015), we considered (i) data from subjects whose performance exceeded chance level exclusively, and (ii) potential effects of stimulus repetition on RTs that have been reported previously (Lewis & Ellis, 2000; Ramon et al., 2011) during estimation of processing speed based on SRTs.

To summarize, in this study we varied the probability of targets’ occurrence to determine the extent to which measured processing speed is influenced by stimulus predictability and repetition. To anticipate our findings, very rapid minSRTs are observed under conditions of high predictability, where the search space is confined to a binary, unambiguous category (gender decision task), or a very small number of target items (familiarity decision with few identities). These findings indicate that “detection of personal familiarity” *per se* requires significantly more time than previously reported, unless “detection” refers to responses towards an extremely restricted number of target identities (Visconti di Oleggio Castello & Gobbini, 2015), which is terminologically more accurately described as target identification (Besson et al., 2017).

## Methods

### Participants

We tested three groups of subjects: the first (n_1_=8; 4 females, mean age: 31, CI [28, 34]) comprised group members from the iBMLab; group members were highly familiar with the target identities, who were their colleagues for several years. The second and third group (n_2_=36; 31 females, mean age: 21, CI [21, 22]; n_3_=27; 24 females, mean age: 22, CI [21, 22]) comprised students from the Department of Psychology who knew the members of Department depicted in the stimulus material through their teaching and mentoring activities. All participants provided written informed consent; all procedures were approved by the internal ethics committee of the Department of Psychology at the University of Fribourg, Switzerland and are in accordance with the Code of Ethics of the World Medical Association (Declaration of Helsinki).

### Stimuli

The full stimulus set comprised natural (uncropped, color) images of 14 facial identities (7 un/familiar) taken from three different viewpoints (frontal, left, right). For each PF identity, images of a corresponding unfamiliar identity carefully matched for age, gender, and appearance (hair color and style, eye color) were taken. Image processing included placement on a uniform grey background (630 × 630 pixels) and correction for low-level properties (luminance, contrast) using the SHINE toolbox (Willenbockel, Sadr, Fiset et al., 2010), as well as additional ones kindly provided by V. Willenbockel to allow for equation of color stimuli.

### Procedures

Prior to completing the experiments, subjects completed familiarity ratings to determine their level of familiarity with each identity (PF and unfamiliar) presented. Each item of the familiarity questionnaire consisted of an image of each stimulus identity taken under varied, natural conditions. Lab members’ images were taken from professional websites, unfamiliar identities’ images were taken from social media. Observers had to indicate their self-reported degree of familiarity with each individual on a scale from 1 (not at all familiar) to 5 (highly familiar). For all experiments stimuli were presented on a 1920 × 1080 pixel VIEWPixx monitor. Subjects’ oculo-motor behavior was recorded at a sampling rate of 1000 Hz with an SR Research Desktop-Mount EyeLink 2K eye tracker (with a chin and forehead rest; average gaze position error ~.5, spatial resolution: ~.01). The eye-tracker had a linear output over the range of the monitor used. Although viewing was binocular, only the left eye was tracked; given the fully balanced stimulus presentation across visual fields, inter-individual differences in ocular dominance were considered irrelevant. The experiment was implemented in Matlab (R2009b, The MathWorks, Natick, MA), using the Psychophysics toolbox (PTB-3) (Kleiner, Brainard, & Pelli, 2007; Pelli, 1997) and EyeLink Toolbox extensions (Cornelissen, Peters, & Palmer, 2002; Kleiner et al., 2007). Calibrations of eye fixations were conducted at the beginning of the experiment using a nine-point fixation procedure as implemented in the EyeLink API (see EyeLink Manual) and using Matlab software. Afterwards, calibrations were validated with the EyeLink software and repeated when necessary until reaching an optimal calibration criterion. Drift correction was performed on each trial via central cross fixation.

In the *gender categorization task*, subjects were instructed to perform choice saccades towards female faces. In this task three images (viewpoint changes) for each PF individual (6 identities, 3 females) and their unfamiliar counterparts were presented. A trial began with a central fixation cross displayed between 800 and 1600ms, followed by a 200ms blank and subsequent presentation of the target/distractor pair presented for 600ms. After a saccade was registered, the next trial was presented after a 1000ms blank inter-trial interval. Stimuli subtended 14˚×14° (average face height was 11°), and stimulus eccentricity was 8.6˚ of visual angle. With all possible combinations and equal number of presentations per identity and visual field, the total number of trials was 216; subjects took self-paced breaks after each block of 54 trials.

The two *familiarity categorization tasks* differed in terms of the number of identities depicted, but both required observers to perform choice saccades towards personally familiar identities presented with unfamiliar distractors. The low and high predictability variants involved presentation of 7 PF (3 females), or 3 PF (all male) identities, respectively, as well as an equal number of well-matched UF distractors. Presentation parameters were identical to those described for the gender categorization task (see above). The procedural parameters paralleled those used by Visconti di Oleggio Castello and Gobbini (2015), with exception of stimulus presentation duration (600ms instead of 400ms), as initial pilot testing revealed slightly longer presentation durations were necessary for acceptable performance levels. On each trial, a PF identity was paired with a same-gender, same-orientation distractor and appeared with equal probability in either visual field. The total number of trials for the low predictability familiarity categorization task amounted to 150. To achieve a comparable number of trials in the high predictability variant for comparison with the low predictability variant, each unique stimulus x visual field combination was presented three times leading to a total 162 trials over three blocks; these trials / blocks were doubled to further determine potential effects of repetition in the high predictability categorization task (see Analyses, section iii.). Subjects took self-paced breaks after each block of 50 or 54 trials (low or high predictability variant), respectively.

## Analyses

### Preprocessing

We applied the adaptive velocity based algorithm developped by Nystrom and Holmqvist (2010) to find the onset of the first saccade (if any) within each trial. We discarded trials in which the onset of the first saccade was lower than 80 ms (Visconti di Oleggio Castello & Gobbini, 2015), as these were considered anticipatory saccades.

### Statistical analyses

As mentioned above, across all experiments we considered only data from subjects whose performance exceeded chance level. For the gender categorization task this led to exclusion of one departmental member (n_1_=7/8); all student participants performed above chance and were considered (n_2_=36/36). For the familiarity categorization tasks, only data from subjects who performed reliably across *both low and high predictability variants* were considered (n_1_=4/8, n_3_=14/27). Analyses performed to determine the effect of stimulus repetition were conducted on data from subjects who performed above chance across all blocks of the *high predictability* familiarity categorization task (n_1_=7/8, n_3_=20/27). Analyses of accuracy and mean SRTs were performed in R (version 3.2.4; R Core Team, 2013) using the lme4 package (Bates, Maechler, Bolker & Walker, 2014) and the lmerTest (Kuznetsova, Brockhoff & Christensen, 2015) to obtain p-values of the fixed predictors of the fitted models. Note that given our research question, we were not interested in between-group differences, but rather those related to stimulus predictability, i.e. those observed *within* groups.

#### i. Gender categorization

Accuracy, mean and minSRTs are reported descriptively for the gender categorization task for all participants (n_1,_ n_2_), as this task served only as a baseline to demonstrate subjects’ ability to perform the SRT task.

#### ii. Personal familiarity categorization: low vs. high stimulus predictability

*Accuracy*. To investigate the effect of stimulus predictability on subjects’ accuracy, we performed generalized linear mixed models with a binomial family (Jaeger, 2008) for the data obtained in the experiments characterized by lower (7 identities), or higher stimulus predictability (3 identities), respectively. This was done separately per group tested given the unequal sample sizes available. In this model, the main predictor is the variable ‘predictability’ (*low* and *high* for larger and smaller number of identities presented) and the variable participant is a random factor. We performed the Log-likelihood Ratio Test to compare the null and full model, and assess the significance of the predictor.

*Mean SRTs*. To investigate the effect of stimulus predictability on SRTs, we performed a linear mixed model for the data obtained in the experiments characterized by lower (7 identities), or higher stimulus predictability (3 identities), respectively, considering only the correct trials.^3^ In this model, the main predictor is the variable ‘predictability’ and the variable participant is a random factor. As for accuracy scores, we performed the Log-likelihood Ratio Test to compare the null and full model, and assess the significance of the predictor.

*Minimum SRTs*. We estimated minSRTs in two different ways^4^. First, across subjects’ trials (i.e., *group minSRTs*), we performed a chi-square test using 10ms time bins across trials. We considered the first bin where the number of correct trials outperforms statistically the number of incorrect trials (*p<.05*), followed by at least 3 significant consecutive bins (Besson et al. 2017). Second, we determined *individuals’ minSRTs*. To this end, following the procedure reported by Besson et al. (2017), we (i) considered only RTs of participants who performed above chance level, and (ii) opted for 40ms time bins using the Fisher’s exact test (*p<.05*). Using this procedure, some participants’ individual minSRT could not be computed (due to proximal in/correct trial distributions). Finally, we assessed the effect of stimulus set size for *individual subjects* of n_3_ using the Wilcoxon signed-rank test (*p<.05*). Note that the comparison of minSRTs as a function of stimulus set size was not conducted for n_1_ due to the insufficient statistical power, as only four participants’ data were considered.

#### iii. Effect of stimulus repetition under conditions of high stimulus predictability

To determine the effect of stimulus repetition, we used the above described procedure for all behavioral measures, however, taking into account *all* correct trials (from *all 6 blocks* of the high predictability familiarity categorization experiment; cf. above). We compared RT associated with the first presentation of a stimulus with each subsequent presentation.

## Results

Table 1 summarizes the results obtained for all subjects for gender categorization, and personally familiar face recognition under low and high predictability conditions, respectively. Individual subjects’ data are reported in Table S 1 and Table S 2; Figure 1 shows individuals’ minSRTs plotted against accuracy scores across all experiments.

**Table 1.** Accuracy (in %) and minimum saccadic reaction times (in ms) obtained across experiments for all subjects tested, whose performance was above chance level

|  | Department members (n<sub>1</sub>) | Students (n<sub>2</sub>) |
| --- | --- | --- |
| Accuracy | 81 [71, 90] | 84 [81, 86] |
| minSRT | 200ms | 140ms |
| Mean SRT | 276 | 252 |
| Median SRT | 262 | 236 |
| CI | [272, 281] | [251, 254] |

***Personal familiarity categorization***
|  | Department members (n<sub>1</sub>) | Students (n<sub>3</sub>) |
| --- | --- | --- |
| **Low predictability** |  |  |
| Accuracy | 74 [64, 81] | 68 [64, 72] |
| minSRT | 360 | 260 |
| Mean SRT | 373 | 334 |
| Median SRT | 390 | 334 |
| CI | [370, 388] | [331, 340] |
| **High predictability** |  |  |
| Accuracy | 82 [74, 89] | 76 [71, 79] |
| minSRT | 260 | 240 |
| Mean SRT | 345 | 345 |
| Median SRT | 353 | 339 |
| CI | [343, 357] | [343, 350] |

95% confidence intervals are provided in brackets.

**Figure 1.**
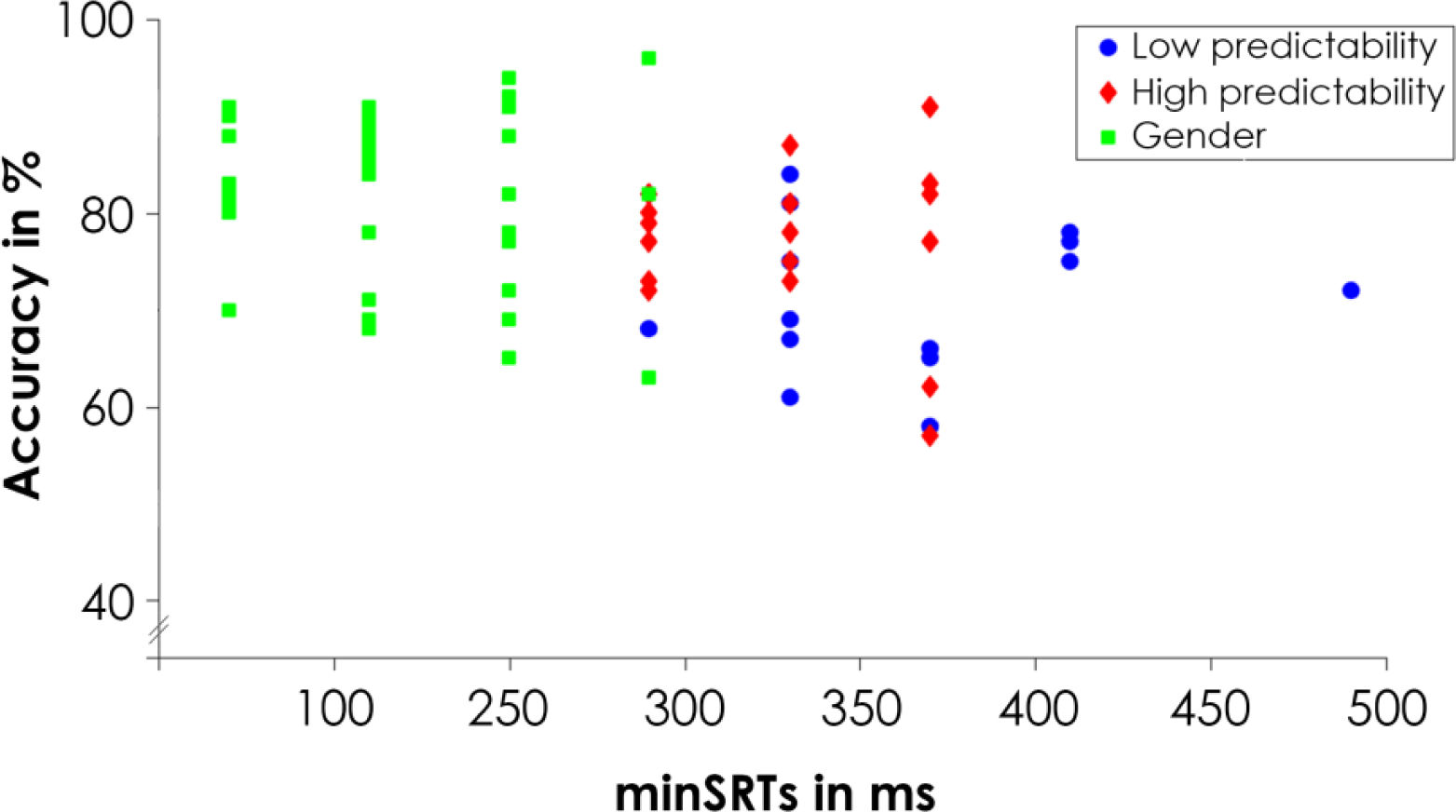
Individual subjects’ minSRTs plotted against performance accuracy. Note that for 3 student subjects minSRTs could not be computed (cf. Table S 2).

### i. Gender categorization

Both departmental members and student participants could reliably perform the gender categorization task, achieving 81% and 84% on average, and exhibiting minimum SRTs of 200ms and 140ms, respectively (see Figure 2).

**Figure 2.**
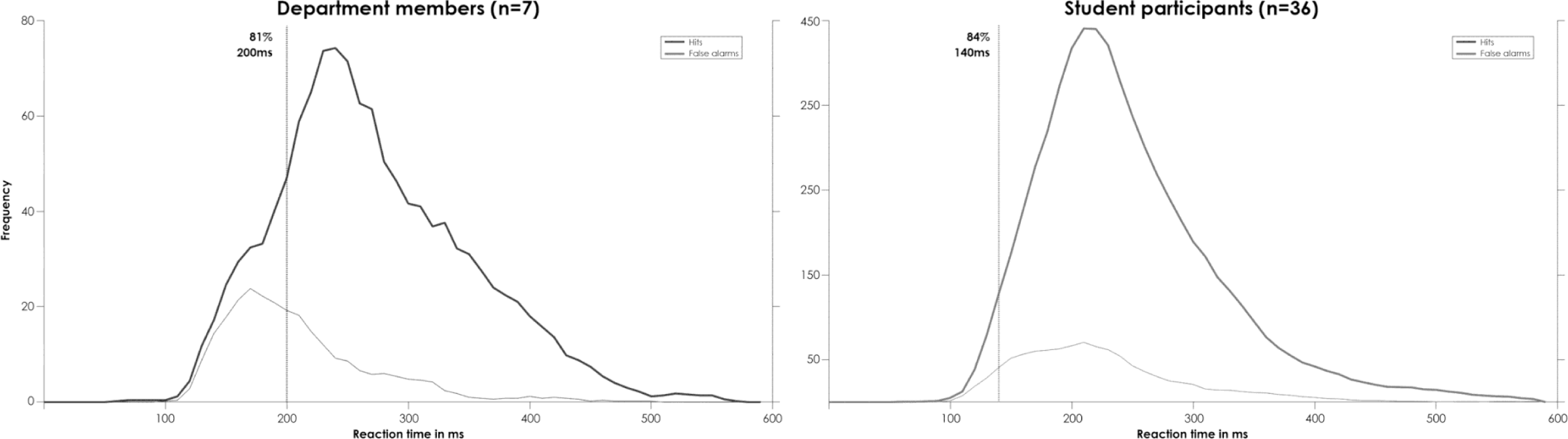
Distributions of participants’ SRTs expressed during the gender categorization task. Hits and false alarms per time bin are indicated as thick and thin lines, respectively. Vertical lines indicate each group’s min SRTs; along with average accuracy.

### ii. Personal familiarity categorization: lower vs. higher stimulus predictability

*Accuracy*. The groups’ SRT distributions for the low and high predictability familiarity categorization tasks are illustrated in Figure 3. For departmental members’ accuracy scores, the fitted model (considering the total of 1248 trials) revealed a significant main effect of stimulus set size (i.e., number of identities depicted; *X*^2^(1)=13.73, *p=.0002*); performance given 7 target identities was significantly lower than when 3 target identities were presented (74% vs. 82%; z=3.69, *p=0.0002*). For students’ accuracy scores, the results from the fitted model (based on a total of 4361 trials) also showed a significant main effect of stimulus set size *X*^2^(1)=33.53, *p<.0001*); again, performance for the lower as compared to higher predictability task variant was significantly lower (68% vs. 76%; z=5.78, *p<.0001*). For parameter estimates of the fixed effects for the generalized linear mixed model with the binomial family for each group tested see Table S 3.

**Figure 3.**
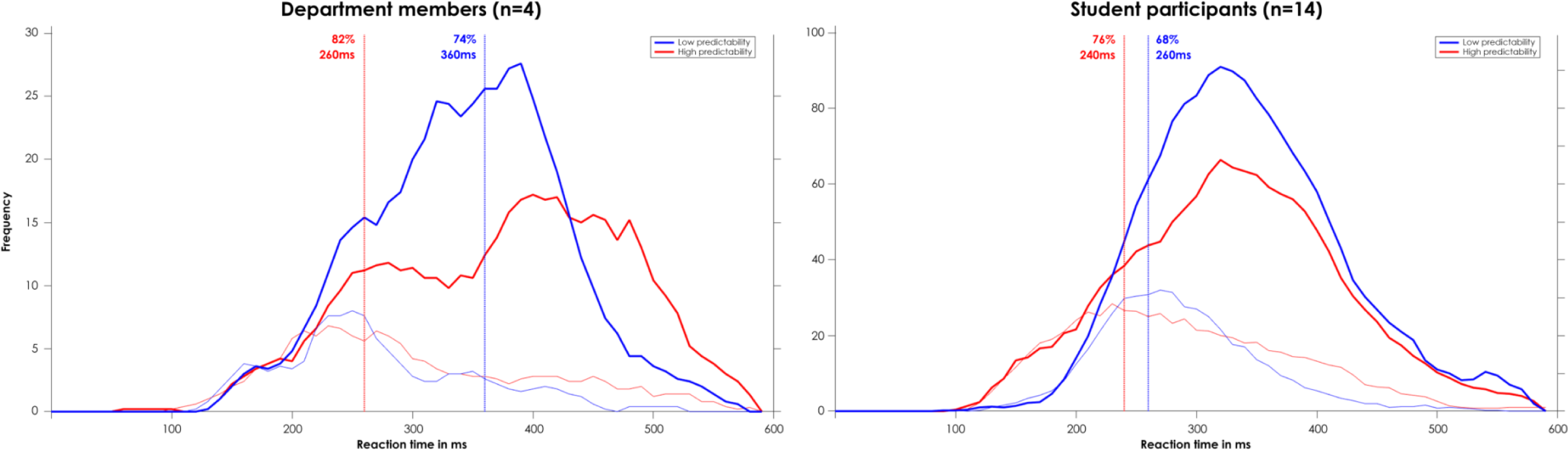
Distributions of participants’ SRTs expressed during the personal familiarity categorization task with low (blue) and high (red) predictability. Hits and false alarms per time bin are indicated as thick and thin lines, respectively

*Mean SRTs*. For departmental members’ SRTs the fitted model (considering the total of 967 trials) revealed a significant main effect of stimulus set size (*X*^2^ (1)=29.70, *p<.0001*). Mean (373 vs 345; t=-5.49, *p<.0001*). For students’ SRTs the results from the fitted model (based on a total of 3142 trials) showed a significant main effect of stimulus set size (*X*^2^(1)=14.67, *p=.0001*). Contrary to the departmental members, the mean of RTs in Experiment 1 was significantly faster than in Experiment 2 (334 vs 345; t=3.83, *p=.0001*). For parameter estimates of the fixed effects for the Linear Mixed Model for each group tested see Table S 3.

*MinSRTs*. The Wilcoxon rank sum test revealed no main effect of stimulus set size on the *individual minSRTS* (see Table S 2) for student participants in n_3_ (*p=.11*).

### iii. Effect of stimulus repetition under conditions of high stimulus predictability

*Accuracy*. The fitted model (considering the total of 1944 trials) revealed no significant effect of repetition on subjects accuracy scores for n_1_ (*X*^2^(5)=2.14, *p=.83*). However a significant main effect of repetition was shown for n_2_ (*X*^2^(5)=16.22, *p<.05*; based on 6480 trials in total). Performance in the first image presentation (73%) was significantly lower than for the fourth image presentation (z=2.61, *p<.05*) and the fifth image presentation (z=3.38, *p<.001*) with a performance of 78% and 79%, respectively (see Table 2). For parameter estimates of the fixed effects for the generalized linear mixed model with the binomial family for each group tested see Table S 5.

**Table 2.** Accuracy, mean and min SRTs for choice saccades towards personally familiar faces as a function of stimulus repetition in Experiment 2 (personal familiarity categorization with high predictability).

| Block | Department members ($n_1$) | | | | Students ($n_2$) | | | |
| --- | --- | --- | --- | --- | --- | --- | --- | --- |
|  | Accuracy<br>in % | Mean<br>SRTs | CI | minSRTs | Accuracy<br>in % | Mean<br>SRTs | CI | minSRTs |
| 1<sup>st</sup> | 77 | 331 | [277; 386] | 250 | 73 | 350 | [325; 374] | 280 |
| 2<sup>nd</sup> | 78 | 307 | [239; 375] | 300 | 74 | 330 | [298;361] | 260 |
| 3<sup>rd</sup> | 75 | 306 | [238; 375] | 340 | 76 | 326 | [294;357] | 240 |
| 4<sup>th</sup> | 76 | 293 | [225; 361] | 290 | 78 | 312 | [280;343] | 220 |
| 5<sup>th</sup> | 74 | 310 | [241; 379] | 260 | 79 | 318 | [286;350] | 220 |
| 6<sup>th</sup> | 74 | 306 | [237; 374] | 290 | 74 | 319 | [287;351] | 240 |

Note that per block each stimulus was shown 3 times.

*Mean SRTs*. The fitted model revealed a significant effect of repetition on mean RTs for n_1_ (*X*^2^(5)=32.18, *p<.0001*) and n_2_ (*X*^2^(5)=126, *p<.0001*) (based on a total 1411 and 4834 trials, respectively). Both groups were slower in the first image presentation compared to the five other image presentations (see Table 2). For parameter estimates of the fixed effects for the Linear Mixed Model for each group tested see Table S 6.

## Discussion

Previous work has demonstrated that the primate brain can visually categorize faces and animals in a highly proficient and rapid manner (e.g. Thorpe et al., 1996; Rousselet et al., 2003; Crouzet et al., 2010; Fabre-Thorpe, 2011; Kirchner & Thorpe, 2006). Visual categorization paradigms, which are considered to involve activation of representations that impact observers’ responses in terms of maximal “presetting” of the visual system for the task at hand (Thorpe et al., 2001), have been deployed to constrain theories of visual processing (for a review see e.g., Fabre-Thorpe, 2011), focusing on the role of stimulus category and information on visual processing.

One body of studies has honed in on visual categorization of a single stimulus category: *faces*. Independent lines of research have reported that both level of processing (e.g., Besson et al., 2017; Barragan-Jason et al., 2012), as well as stimulus familiarity (e.g., Visconti di Oleggio Castello & Gobbini, 2015; Ramon et al., 2011; Ramon & Gobbini, 2018) can affect overtly observed face processing proficiency. Besson et al. (2017) reported that *manual* minRTs increased systematically as a function of task performed, with higher proficiency exhibited for human face (vs. animal) categorization (~240ms) and search for a predefined target identity (“individual face recognition”, ~260ms). Famous face recognition on the other hand, i.e. deciding whether a given identity of *many possible* ones was famous or not, took substantially longer (~380ms). Collectively, these findings suggest that detecting the presence of a face precedes determining its familiarity, or identity (Besson et al., 2017; Barragan-Jason et al., 2012).

Other behavioral findings, however, challenge this view. Visconti di Oleggio Castello and Gobbini (2015) recently reported that observers could perform SRTs towards personally familiar faces in as little as 180ms; even faster SRTs to famous face images were reported by Mathey et al. (2014; <140ms). Based on their findings, Visconti di Oleggio Castello and Gobbini (2015) suggested that processing of familiarity is rapidly detected at this latency, “prior to explicit recognition of identity”. This notion is incompatible with the earliest reported neural markers of familiarity recognition (>170ms; Barragan-Jason et al., 2015; Caharel et al., 2014; Huang et al., 2017).

In the present study we reasoned that these apparently conflicting findings can be reconciled by careful consideration of the effect that both task demands and expectations exert on visual categorization, as well as the terminology adopted to communicate findings (Ramon, in press). Specifically, we proposed that procedural aspects effectively constrain the decisional space within which observers perform a task, which in turn determines measured behavior (Ramon, in press; Ramon et al., in press; see also Ramon & Rossion, 2010; Ramon, Busigny, Gosselin & Rossion, 2017; Ruffieux, Ramon, Lao, Colombo, Stacchi et al., 2017). To test this assumption, we measured observers’ saccadic RTs across three experiments. Each experiment required observers to perform binary decisions, expressed through choice saccades towards a predefined target category. These target categories varied across tasks: female faces in the gender decision task, personally familiar face in the familiarity decision tasks. Importantly, we deployed two familiarity decision tasks, which differed in terms of the total number of possible target identities (3 vs. 7), thereby manipulating stimulus predictability, and hence breadth of decisional space. Importantly, all observers were presented the *exact same target identities* (i.e., stimuli considered as PF and UF faces), and we determined potential effects of stimulus repetition on processing speed.

The findings obtained for the *gender categorization* task reveal that visual processing can be extremely efficient and fast when operating within a narrow decisional space. Student participants exhibited high performance accuracy (84%) and very rapid minSRTs when performing choice saccades towards female faces (~140ms). Department members exhibited slightly lower accuracy (81%) and somewhat longer minSRTs (200ms), which we attribute to the smaller hit-to-false alarm ratio given the smaller sample size (6 vs. 36 participants) and hence number of data points available. In the context of *familiarity categorization* tasks, we found that participants’ performance was modulated by the number of PF identities presented. Accuracy increased and RTs decreased with fewer target identities presented (7 vs. 3 PF identities), and high target predictability was associated with minSRTs of ~250ms.

### Reconciling discrepancies in minSRTs reported for personally familiar face categorization

We believe that procedural differences may at least partially account for the difference in minSRTs for familiarity decisions observed here and elsewhere (180ms; Visconti di Oleggio Castello & Gobbini, 2015). First, following others’ adopted procedures (e.g., Besson et al., 2017; Ramon et al., 2011; Ramon et al., 2017b; Ramon et al., in press) we included only data of subjects who exhibited reliably high performance, as opposed to also including correct trials from individuals who were at or near chance level (cf. Visconti di Oleggio Castello & Gobbini, 2015). Although we did not find a systematic relationship between accuracy and minSRTs among the subjects considered for our analyses (see Figure 1), we note that lab members’ RTs decreased by 100ms between the low and high predictability familiarity decision experiments. In comparison, students achieved lower accuracy in general (68% and 76%) relative to lab members (73% and 82%), and across tasks did not exhibit the decrease in minSRTs observed for lab members. That is, students performed at maximum speed during the low predictability experiment, but could increase their performance *accuracy* due to a narrowing of the decisional space given less target identities. Regardless of whether groups differed regarding which behavioral measure improved by predictability, we believe that the comparable minSRTs exhibited across groups (lab members: 260ms, students: 240ms) reveals the fastest processing speed for a 1-of-three target identity categorization scenario when estimates are based on highly reliable performance.

Moreover, in our study PF and UF stimuli were identical to all subjects (cf. e.g., Besson et al., 2017; Mathey et al., 2012; Ramon et al., 2011; Ramon et al., 2017a, b; Caharel et al., 2015). This is fundamentally different from Visconti di Oleggio Castello and Gobbini’s (2015) study, in which one subjects’ PF stimuli served as UF stimuli for another. We suggest that the rapid minSRTs of 180ms these authors reported are thus more likely to reflect effective top-down “presetting” towards diagnostic features of specific images, which can lead to a significant decrease in minSRTs (Delorme, Rousselet, Macé & Fabre-Thorpe, 2004). Therefore, we suggest that not necessarily the number of distractor items, but rather the *overall number of potential targets to verify determines visual processing speed in familiarity decisions*.

The repetition effects reported here and elsewhere using stimulus set sizes ranging from one famous (Lewis & Ellis, 2000), to 27 PF identities (Ramon et al., 2011) lend further support to the idea that RTs decrease due to activation of image features; previously observed task-learning effects (cf. Visconti di Oleggio Castello & Gobbini, 2015) can also further contribute to a systematic decrease in recorded RTs. Additional support for this claim stems from SRT differences between studies that have both employed time sensitive go/no-go paradigms. Ramon et al. (2011) reported minSRTs and mean SRTs of 380ms and 510ms. We attribute the shorter manual response latencies compared to those reported by Barragon-Jason et al. (2012: 467ms and 581ms; 2015: 460ms and 635ms) to the difference in stimulus set size and hence target predictability. Collectively, our findings stress the importance of considering the task’s *decisional space* and *stimulus repetition*, as both *significantly affect the speed of familiarity categorizations*.

### A formal framework of saccadic reaction times to face stimuli

In the present study, we focused on the impact of stimulus probability, or decisional space, and stimulus repetition as determinants of subjects’ minimum processing speed during visual categorization. However, an additional factor which can further modulate expressed behavior, which we did not systematically control here, is *visual information*. This point can be demonstrated by considering for example visual categorization of facial expression. Similar to the present study, the number of categories (i.e. facial expressions) presented will determine the breadth of decisional space, and thus likely modulate subjects’ performance. Moreover, however, the *type of information* based on which categorical decisions are performed will also play an important role. Inclusion of facial expressions that are more likely confused by subjects (e.g., Jack, Blais C, Scheepers, Schyns & Caldara, 2009; Jack, Garrod, Yu, Caldara & Schyns, 2012; Rodger, Vizioli, Ouyang & Caldara, 2015) will lead to an increase in task difficulty and hence decrease in accuracy and/or delayed RTs.

Beyond the *objectively available* visual information, inter-individual anatomical differences that may affect information processing efficiency would also require consideration. One could speculate that anatomical differences in early visual cortices (cortical thickness, surface area) reported to affect perceptual discrimination and mental imagery (Bergmann, Genc, Kohler, Singer & Pearson, 2016; Song, Schwarzkopf, Kanai & Rees, 2015) may affect processing of the parafoveally presented visual information subject to categorical responses in SRT paradigms. A comprehensive account of processing speed based on which a predictive model of saccadic choices for a particular task can be elaborated will require consideration of stimulus probability, as well as visual information and anatomically constrained information processing proficiency.

### Implications beyond visual categorization measured behaviorally: neuroimaging of face recognition

We suggest that effects of stimulus predictability may also account for seemingly contradictory neuroimaging findings reported previously. Electrophysiological studies determining the influence of familiarity have consistently reported familiarity-dependent modulation of later (>400ms) components, while effects on earlier (<200ms) components are more variable (for reviews see Huang, et al., 2017; Ramon & Gobbini, 2018). For example, some studies suggest that the N170 / M170 – the earliest face-sensitive component electrical signal recorded from the scalp (Bentin & Deouell, 2000; Eimer, 2000) is sensitive to familiarity and task (Rossion et al., 1999) –. These studies have, however, reported enhanced (Caharel, Courtay, Bernard, Lalonde & Rebai, 2005; Caharel, Fiori, Bernard, Lalonde & Rebai, 2006; Kloth, Dobel, Schweinberger, Zwitserlood, Bolte et al., 2006; Wild-Wall, Dimigen & Sommer, 2008), as well as decreased N170 amplitude (Todd, Lewis, Meusel & Zelazo, 2008). Of note is that active, identity-related tasks generally lead to modulation of early components, compared to passive viewing or orthogonal tasks, for which later familiarity effects have been more consistently reported (Huang et al., 2017; Taylor, Shezad & McCarthy, 2016).

As reviewed by Ramon and Gobbini (2018), whether or not familiarity modulation can be found for early electrophysiological components may depend on pre-activation of associative knowledge. Studies reporting early modulation for PF faces have in common that participants usually know in advance the identities presented (e.g. Caharel et al., 2014). Similarly, studies reporting early modulation for famous faces often involve stimulus selection based on subjects’ previously assessed level of familiarity with various famous individuals (Huang et al., 2017). Lastly, studies using experimentally learned faces have also reported familiarity-dependent modulation of early components (Taylor et al., 2016). In all of these studies pre-activation of identity information can be regarded as defining the decisional space from which to-be-presented identities are drawn. Based on our findings, and reports of expectation-related N170 modulation (e.g. Johnston, Overell, Kaufman, Robinson & Young, 2016), we hypothesize that the likelihood of observing familiarity-dependent modulation of early electrophysiological components, and neural responses measured with functional magnetic neuroimaging (Ramon & Gobbini, 2018), will increase with predictability of familiar exemplars presented.

### Conclusions

Based on the present and previous findings we emphasize the importance of adopting a terminology that takes into consideration more than merely the *stimulus* being processed. The *process* involved, which depends on the opted-for paradigm, crucially impacts processing speed (cf. Besson et al., 2017). For instance, the term “face recognition” or “detection” is oftentimes used across a range of tasks and without sufficient consideration of differences in specific procedures implemented, leading to at times seemingly contradictory conclusions (Besson et al., 2017; Ramon, in press). Our findings that target predictability and repetition affect processing speed in binary forced-choice paradigms emphasize the importance of simultaneously considering task constraints and procedural aspects when attempting to determine processing speed to constrain theories of visual processing. At the very least we suggest that previous findings of SRT modulation attributed to personal familiarity should be revisited: the minSRTs reported by Visconti di Oleggio Castello and Gobbini (2015) do not reflect facilitated familiar face detection per se, but rather demonstrate how powerful top-down predictions can expedite rapid visual orienting towards specific, *expected* images. These findings also provide a basis to account for seemingly contradicting findings regarding familiarity-related modulation of responses to faces recorded with neuroimaging techniques, which are likely affected by the operational decisional space and “presetting” due to subjects’ expectations.

## Acknowledgements

We thank our previous Bachelor students Nour Al-Khodairy, Morgane Bulliard, Aurelie Christinaz, and Lysiane Constantin for their invaluable help during data collection.

## Supplementary Material

**Table S 1.** Subjects’ accuracy (in %), mean and minimum saccadic reaction times (in ms, with 95 confidence intervals) obtained for the gender categorization task.

|  | Accuracy | minSRT | Mean SRT |
| --- | --- | --- | --- |
| **Department members (n<sub>1</sub>)** |  |  |  |
| 1 | 92 | 200 | 279 [272; 288] |
| 2 | 94 | 200 | 250 [244; 255] |
| 3 | 65 | 200 | 177 [170; 183] |
| 4 | 82 | 240 | 344 [330; 358] |
| 5 | 78 | 200 | 248 [239; 257] |
| 6 | 96 | 240 | 346 [338; 354] |
| 7 | 63 | 240 | 237 [229; 244] |
| **Students (n<sub>2</sub>)** |  |  |  |
| 1 | 91 | 200 | 286 [280; 294] |
| 2 | 89 | 160 | 264 [256; 273] |
| 3 | 82 | 120 | 174 [168; 179] |
| 4 | 88 | 120 | 205 [199; 212] |
| 5 | 86 | 160 | 233 [225; 241] |
| 6 | 90 | 160 | 328 [315; 341] |
| 7 | 70 | 120 | 176 [170; 185] |
| 8 | 78 | 160 | 188 [181; 196] |
| 9 | 68 | 160 | 224 [215; 232] |
| 10 | 90 | 160 | 231 [224; 238] |
| 11 | 91 | 160 | 253 [245; 263] |
| 12 | 88 | 160 | 230 [224; 235] |
| 13 | 82 | 200 | 381 [364; 399] |
| 14 | 87 | 160 | 267 [257; 279] |
| 15 | 91 | 120 | 216 [210; 224] |
| 16 | 85 | 160 | 251 [241; 264] |
| 17 | 71 | 160 | 282 [267; 298] |
| 18 | 84 | 160 | 197 [193; 202] |
| 19 | 81 | 120 | 254 [243; 265] |
| 20 | 83 | 120 | 297 [281; 312] |
| 21 | 72 | 200 | 268 [256; 281] |
| 22 | 89 | 160 | 226 [220; 232] |
| 23 | 88 | 200 | 272 [265; 279] |
| 24 | 88 | 120 | 178 [173; 182] |
| 25 | 91 | 160 | 202 [195; 208] |
| 26 | 89 | 120 | 232 [224; 239] |
| 27 | 82 | 120 | 224 [220; 229] |
| 28 | 88 | 200 | 281 [274; 288] |
| 29 | 86 | 200 | 275 [270; 281] |
| 30 | 90 | 160 | 310 [297; 323] |
| 31 | 70 | 200 | 319 [307; 333] |
| 32 | 78 | 160 | 260 [252; 268] |
| 33 | 68 | 200 | 266 [257; 274] |
| 34 | 90 | 200 | 250 [245; 256] |
| 35 | 91 | 160 | 237 [232; 243] |
| 36 | 88 | 160 | 290 [282; 298] |

**Table S 2.**
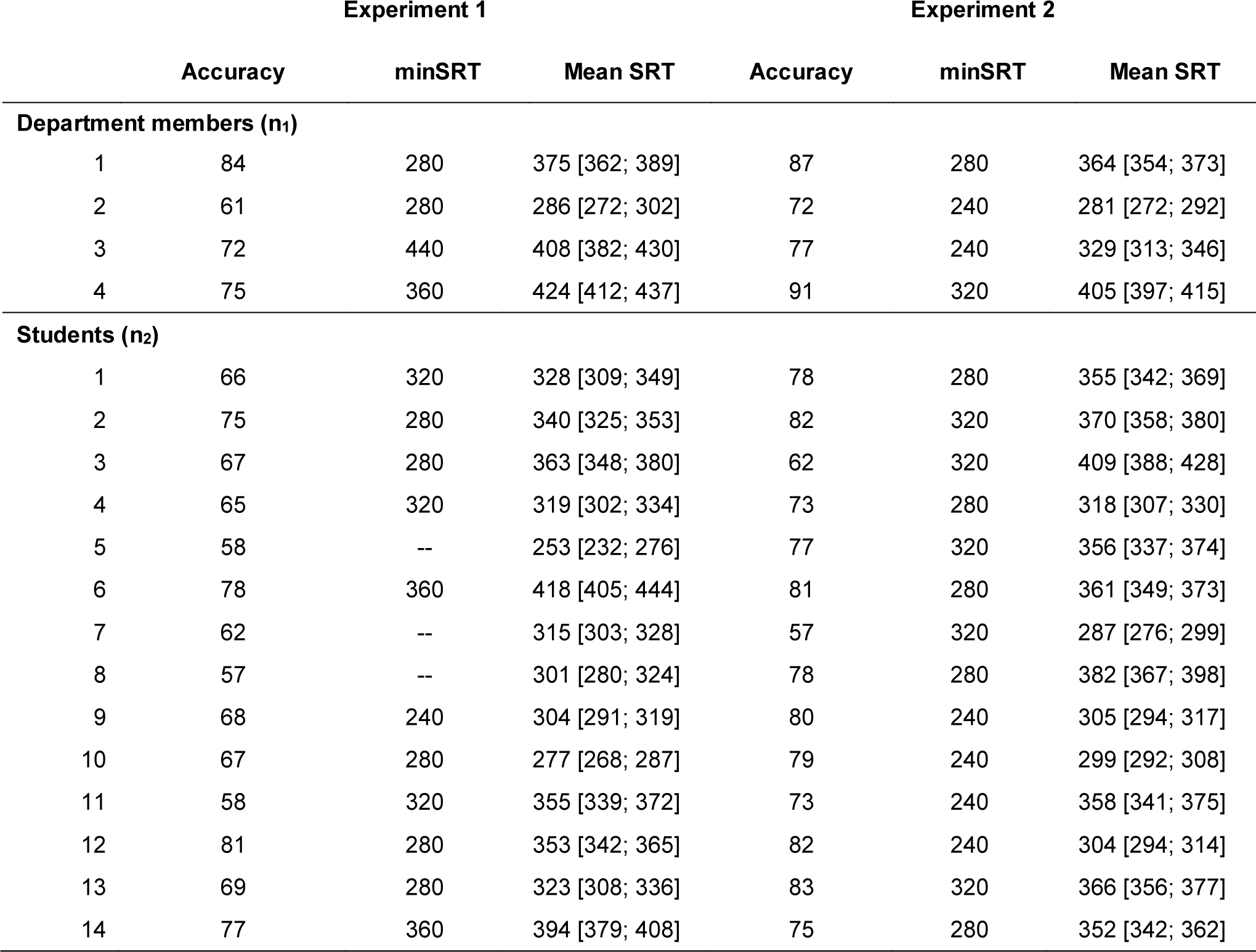
Subjects’ accuracy (in %), mean and minimum saccadic reaction times (in ms, with 95 confidence intervals) for Experiment 1 and Experiment 2.

**Table S 3.** Parameter estimates of the fixed effects for the generalized linear mixed model with the binomial family on accuracy

| Fixed Effects | log(Odds) | SE | CI | Odds | Odds CI | z-value | p-value |
| --- | --- | --- | --- | --- | --- | --- | --- |
| **Lab Members** |  |  |  |  |  |  |  |
| Intercept | 1.03 | .22 | [.48; 1.6] | 2.80 | [1.61 4.95] | 4.57 | <.0001 |
| high predictability | .51 | .13 | [.24; 0.78] | 1.66 | [1.27; 2.19] | 3.69 | .0002 |
| **Students** |  |  |  |  |  |  |  |
| Intercept | .77 | .09 | [.59; .94] | 2.16 | [1.80; 2.58 ] | 8.74 | <.0001 |
| high predictability | .40 | .07 | [.26; .53] | 1.49 | [1.30; 1.70] | 5.78 | <.0001 |

**Table S 4.**
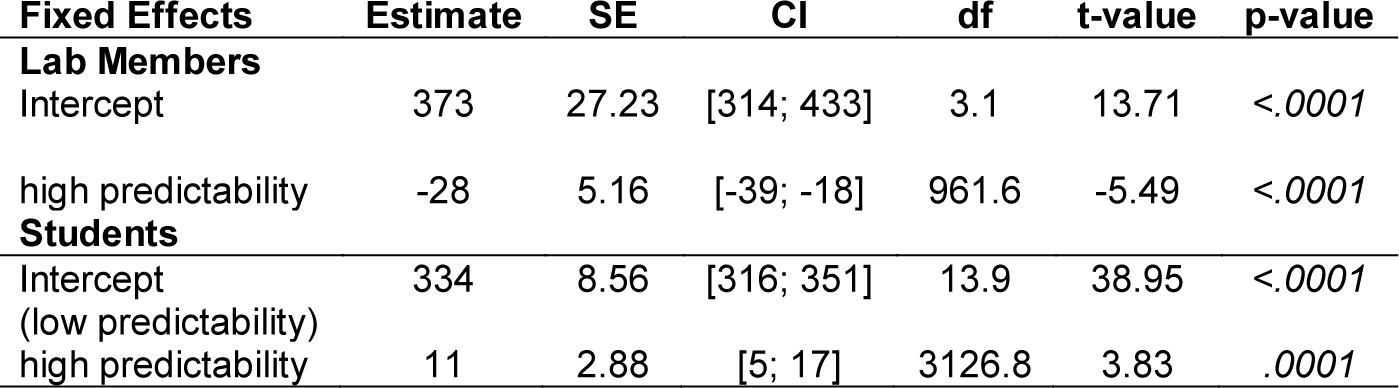
Parameter estimates of the fixed effects for the Linear Mixed-Effects Model on SRTs.

**Table S 5.** Parameter estimates of the fixed effects for the generalized linear mixed model with the binomial familyl on accuracy

| **Fixed Effects** | **log(Odds)** | **SE** | **CI** | **Odds** | **Odds CI** | **z-value** | **p-value** |
| --- | --- | --- | --- | --- | --- | --- | --- |
| **Lab Members** |  |  |  |  |  |  |  |
| Intercept | 1.21 | .37 | [.37; 2.06] | 3.35 | [1.44; 7.84] | 3.24 | .001 |
| Block2 | .05 | .19 | [-.32; .42] | 1.05 | [.73; 1.53] | .28 | .77 |
| Block3 | -.10 | .18 | [-.47; .26] | .90 | [.63; 1.30] | -.56 | .58 |
| Block4 | -.03 | .19 | [-.40; .33] | .97 | [.67; 1.39] | -.19 | .85 |
| Block5 | -.13 | .18 | [-.50; .22] | .88 | [.61; 1.25] | -.74 | .46 |
| Block6 | -.17 | .18 | [-.53; .19] | .84 | [.59; 1.21] | -.92 | .36 |
| **Students** |  |  |  |  |  |  |  |
| Intercept | .97 | .12 | [.73; 1.22] | 2.64 | [2.07; 3.39] | 7.98 | <.0001 |
| Block2 | .06 | .10 | [-.13; .25] | 1.06 | [.88; 1.29] | .64 | .52 |
| Block3 | .16 | .10 | [-.03; .36] | 1.17 | [.97; 1.43] | 1.64 | .10 |
| Block4 | .26 | .10 | [.07; .46] | 1.30 | [1.07; 1.58] | 2.61 | <.05 |
| Block5 | .34 | .10 | [.15; .54] | 1.40 | [1.16; 1.72] | 3.38 | <.01 |
| Block6 | .09 | .10 | [-.10; .29] | 1.10 | [.90; 1.34] | .94 | .35 |

**Table S 6.** Parameter estimates of the fixed effects for the Linear Mixed-Effects Model on SRTs.

| **Fixed Effects** | **Estimate** | **SE** | **CI** | **df** | **t-value** | **p-value** |
| --- | --- | --- | --- | --- | --- | --- |
| **Lab Members** |  |  |  |  |  |  |
| Intercept | 331 | 26 | [277; 386] | 5 | 12.81 | <.0001 |
| Block2 | -24 | 6.94 | [-38; -11] | 1400 | -3.49 | <.0001 |
| Block3 | -25 | 7 | [-39; -11] | 1400 | -3.55 | <.0001 |
| Block4 | -39 | 6.98 | [-52; -25] | 1400 | -3.55 | <.0001 |
| Block5 | -21 | 7.03 | [-35; -7.2] | 1400 | -2.99 | .002 |
| Block6 | -26 | 7.04 | [-39; -12] | 1400 | -3.64 | .0002 |
| **Students** |  |  |  |  |  |  |
| Intercept | 350 | 12.27 | [325; 374] | 21 | 28.51 | <.0001 |
| Block2 | -20 | 3.74 | [-27; -13] | 4807 | -5.31 | <.0001 |
| Block3 | -24 | 3.72 | [-31; -17] | 4807 | -6.50 | <.0001 |
| Block4 | -38 | 3.70 | [-45; 31] | 4807 | -10.22 | <.0001 |
| Block5 | -32 | 3.68 | [-39; -24] | 4807 | -8.58 | <.0001 |
| Block6 | -30 | 3.73 | [-38; -23] | 4807 | -8.15 | <.0001 |

## Footnotes

1 Note that observers responded to different target stimuli (identities personally familiar to one subject served as unfamiliar faces to others), and therefore different visual information. The authors state that “[f]or three subjects one of the target familiar faces had a darker skin color than the other faces. We rejected all the trials containing images of this individual (both as a target and as a distractor) to avoid any bias due to a skin color difference.” (Visconti di Oleggio Castello & Gobbini, 2015; p.5). In our opinion, undesired effects of input-dependent biases *cannot* be abolished through exclusion of individual responses.

2 To our knowledge, Visconti di Oleggio Castello and Gobbini’s (2015) study is the only one to compute SRTs based on correct responses from *all* subjects – including those who performed at/near chance level for choice saccades directed towards PF faces (see Table S6, Visconti di Oleggio Castello & Gobbini, 2015). Min(S)RT estimation in the aforementioned studies involved exclusion of data from observers, who were unable to perform the task (e.g. Besson et al., 2017; Ramon et al., 2011; Ramon et al., 2017; Ramon et al., in press).

3 To account for the different number of trials across experiments, note that we considered *all* trials (150) from the low, and those of first 3 blocks (162) for the low and high predictability variants of the familiarity categorization tasks.

4 Here, as for mean SRTs, only the first 3 blocks were considered.

